# Diversity and characteristics of the oral microbiome influenced by race and ethnicity

**DOI:** 10.1101/2024.10.07.617037

**Authors:** Qingguo Wang, Bing-Yan Wang, She’Neka Williams, Hua Xie

## Abstract

Periodontitis disproportionately affects racial/ethnic populations. Besides social determinants contributing to disparities in periodontal health, variations of oral microbial communities may also be a key factor influencing oral immune responses. To characterize the oral microbiome from different racial/ethnic populations, we collected 161 dental plaque samples from African Americans (AAs), Caucasian Americans (CAs), and Hispanic Americans (HAs) with clinical gingival health or biofilm-induced gingivitis on an intact periodontium. Using metagenomic sequencing, we found significant difference in diversity and abundance of microbial taxa in the dental plaque samples from AA, CA, and HA groups and unique microbial species that can only be detected in a particular racial/ethnic group. Moreover, we revealed racial/ethnic associated variations in functional potential of the oral microbiome, showing that diversity and abundance of antibiotic resistant genes were greater in the oral microbiome of the AAs than those in CAs or HAs, and that the AAs exhibited higher levels of genes involving in modification of glycoconjugates, oligo- and polysaccharides. These findings indicate more complex and higher virulence potential oral microbiome in AA and HA populations, which likely contributes to higher prevalence of periodontitis in AAs and HAs.

**Importance:** Recognizing the variations in the oral microbiome among racial/ethnic populations offers insight into the microbial determinants contributing to oral health disparities. In the study presented here, we found a higher level of bleeding on probing (BOP), an indicator of tissue inflammatory response, in the AA group, which is correspondence with a more complex oral microbiome detected in this group. Our observations suggest that the variations of the oral microbiome associated with racial/ethnic backgrounds may directly relate to their virulence potential including their abilities to induce host immune responses and to resist antibiotic treatment. Therefore, these finding can be a stepping stone for developing precision medicine and personalized periodontal prevention/treatment and for reducing oral health disparities.

## INTRODUCTION

Periodontitis is a prevalent human disease affecting approximately 42% of adults aged 30 and older in the US. The National Health and Nutrition Examination Survey (2009-2014) highlighted significant racial and ethnic disparities in periodontitis, with higher rates observed in African Americans (AAs, 56.6%) and Hispanic Americans (HAs, 59.7%) compared to Caucasian Americans (CAs, 37.0%)(1). These disparities are influenced by multiple factors, including health care access and socioeconomic status (SES) (2, 3). A growing body of evidence shows that human microbiome are dynamic microbial communities that shift with states of health (4). Microbiome dysbiosis has been identified in a long list of diseases using meta-omic techniques, including increased *Staphylococcus aureus* in the skin of atopic dermatitis (5) and decreased diversities and increased Fusobacteria in the gut microbiome in patients with colorectal cancer (6). Interindividual differences of microbiome are also influenced by genetics, environment, social determinants, and lifestyles (7, 8), which leads to distinctive susceptible of population with different racial/ethnic backgrounds to diseases. A recent study demonstrated a critical developmental window of gut microbiome variation at or shortly after 3 months of age, which is driven by social status and environments and may lead to health disparities in adult (9). While oral microbial dysbiosis is a recognized factor in oral diseases (10, 11), there remains a gap in understanding how variations in the oral microbiome contribute to the disparities. Recent studies reported a differential oral microbiota in the populations with oral malignant disorders and oral cancer when compared to that found in the healthy counterparts (12–14). Although signatures of the oral microbiome associated with oral cancer has not yet established, some bacterial taxa, including *Fusobacterium*, *Prevotella*, and *Porphyromonas*, were more abundant in the oral cavities of patients with oral cancer. Population-level analysis of oral microbiome variation was also investigated in racial and ethnic groups. A study by Yang et al. found a significant racial difference in oral microbiome between African ancestry and European ancestry (15). The study of deep sequencing of 16S rRNA genes revealed significantly increased abundances of four periodontitis-associated bacteria, including *Porphyromonas gingivalis*, *Prevotella intermedia*, *Treponema denticola*, and *Filifactor alocis*, in mouth rinse samples of African ancestry, compared to those in the samples from European ancestry, although periodontal status of the cohort in this study is not known.

We previously investigated microbiologic risk factors associated with periodontal health disparities using qPCR. we determined and compared several key members of oral bacteria that play distinct roles in periodontal health in AAs, CAs, and HAs. We detected much higher levels of *P. gingivalis* in the samples from the HA and AA periodontitis patients than in samples from the CA patients, which appears to link to higher index of bleeding on probing observed in the HAs and AAs with periodontitis (16). In addition, we examined *P. gingivalis* level in the oral cavities of intact periodontium individuals with different racial/ethnic backgrounds. Although significant difference of *P. gingivalis* distributions in these groups was not found using qPCR in our previous study, the levels of *P. gingivalis* were higher in the HAs than that in the CAs (17). To further identify population associated differential microbial profiles, we selected a total of 161 dental plaque samples from AAs, CAs and HAs with intact periodontium for whole metagenome shotgun sequencing. We identified significant difference in numbers of non-redundant bacterial genes and diversity and abundance of microbial species among AA, CA, and HA groups. Additionally, several bacterial species were unique in a particular racial/ethnic group. Moreover, functional potentials of the oral microbiome, such as antibiotic resistance and LPS production, were higher in AA group than CA and HA groups. These results suggest that racial/ethnic specific oral microbiome attributes to oral health disparities.

## RESULTS

### Baseline characteristics of cohort

In the previous studies, we determined detection rates of *P. gingivalis*, using qPCR, among 340 individuals with intact periodontium with different racial/ethnic backgrounds (17). Detection rates of *P. gingivalis* were 21.2% of the AAs, 18.2% of the CAs, and 30.6% of the HAs, respectively. To further investigate variations of the oral microbiome associated with periodontal healthy disparities, we selected dental plaque samples with *P. gingivalis* from the cohort and paired them with the *P. gingivalis* negative samples with matched age, gender, and racial/ethnic background, resulting in 44 AAs (GpgA), 42 CAs (GpgC) and 75 HAs (GpgH) (Table 1). GpgA, GpgC, and GpgH were further divided based on absence (Gpg1A, Gpg1C, and Gpg1H) or presence (Group2A, Gpg2C, and Gpg2H) of *P. gingivalis* determined previously by qPCR (17). Statistically significant differences were found in age and gender between AA and HA and between CA and HA groups, but not between AA and CA (Table 1). Higher levels of BOP were found in AAs, compared to CAs and HAs (Negative Binomial Regression, *p* = 0.003 after adjusted for covariates including age and gender.

**Table 1.** Characteristics of the study cohort.

| Characteristics | AAs | CAs | HAs | *p*-value |
| --- | --- | --- | --- | --- |
| Gender (Male/Female) | 20/24 | 16/26 | 17/58 | 0.027<sup>+</sup> |
| Age (year; Mean ± SD) | 44.84±2.84 | 48.63±2.85 | 41.08±12.58 | 0.029<sup>#</sup> |
| BOP (%; Mean ± SD)<sup>a</sup> | 34.92±5.23 | 17.30±3.11 | 22.53±18.29 | 0.006* |
| PI (%; Mean ± SD)<sup>b</sup> | 61.04±5.87 | 50.12±5.24 | 44.94±25.10 | 0.093 |
| Tooth number (Mean ± SD)<sup>c</sup> | 26.72±0.47 | 27.04±0.35 | 27.17±2.22 | 0.963 |

<sup>a</sup> BOP: Bleeding on probing
<sup>b</sup> PI: Modified O'Leary plaque index
<sup>c</sup> Tooth number is based on a total of 32 teeth.
<sup>+</sup> The gender distribution of AA is significantly different from that of HA and there is no significant difference between AA and CA or between CA and HA (Chi-Square Test).
<sup>#</sup> The age distribution of AA is significantly different from other two groups which do not have significant difference in ages (Kruskal-Wallis test).
\* The BOP of AA is significantly different from those of CA and HA which do not differ significantly in BOP (Negative Binomial Regression, adjusted for covariates Age and Gender).

### Diversity and similarity of the oral microbiota among AA, CA, and HA groups

A total of 2,025,809 non-redundant genes were predicted from 161 dental plaque samples using MetaGeneMark. As shown in Fig. 1 and Table 2, the median number of non-redundant genes per sample in AA groups were significantly higher than those found in their counterparts CAs and HAs (Negative Binomial Regression, *p* < 0.001) after adjusting for age and gender. There was no significant difference in number of non-redundant genes between CAs and HAs, regardless of *P. gingivalis* detection with qPCR. In addition, unique genes were detected in each racial/ethnic group. We found the highest number of unique genes (90,477) in the AA group followed by those (48,215 and 27,074) in HAs and CAs respectively (Fig. 1B). These finding suggest a more complex microbial composition in AA population compared to HAs and CAs, which was also confirmed by taxonomic diversities among these groups.

**Fig. 1.**
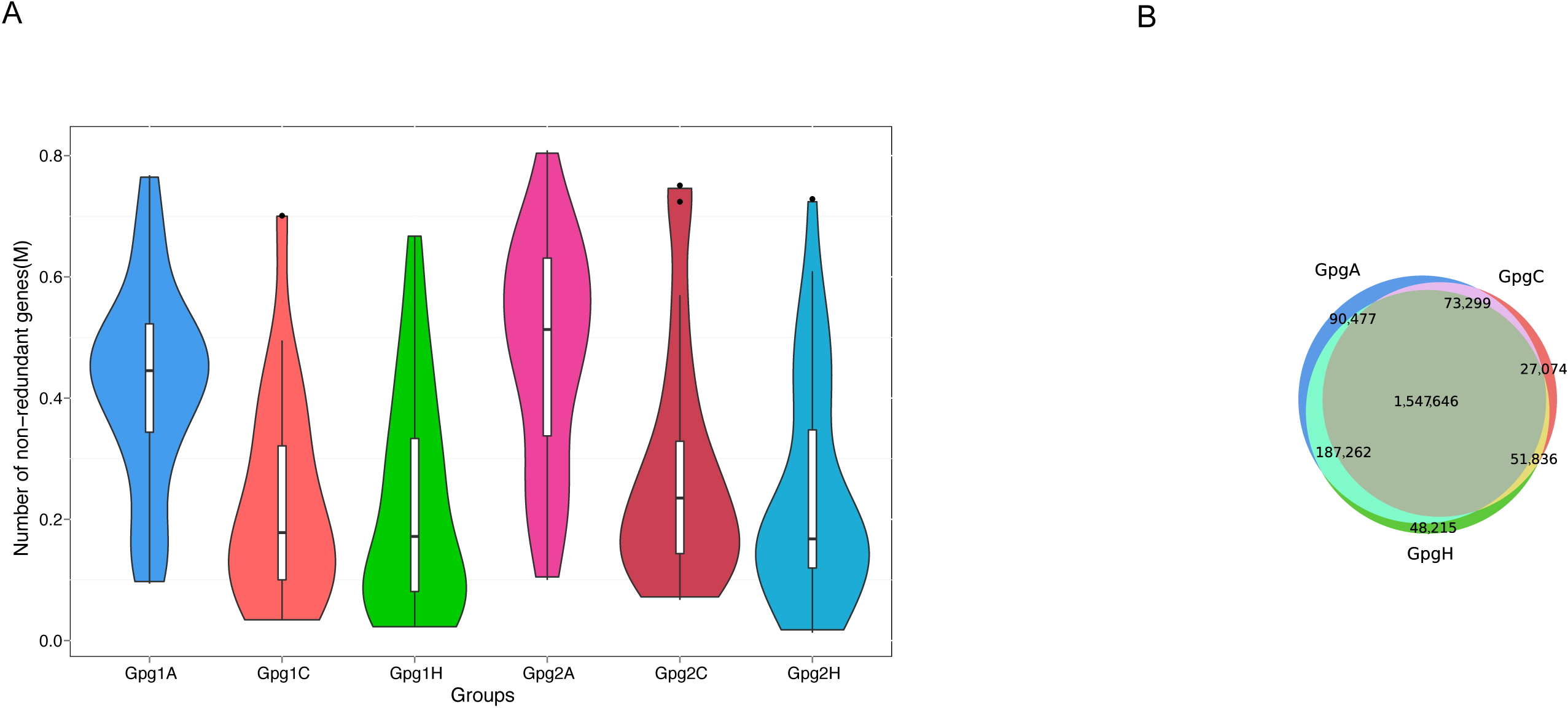
Comparison of the number of non-redundant genes in dental plaque samples from different racial/ethnic groups. (A) The violins represent the richness of non-redundant genes in the samples with low levels of *P. gingivalis* in AAs (Gpg1A), CAs (Gpg1C), and HAs (Gpg1H) or high levels of *P. gingivalis* in AAs (Gpg2A), CAs (Gpg2C), and HAs (Gpg2H). (B) Venn diagram of the total number of non-redundant genes identified in the dental plaques samples from AAs (GpgA), CAs (GpgC), and (GpgH).

**Table 2.** Non-redundant genes identified per samples in different groups.

| Comparison of<br>sample groups | Median number of<br>genes per sample | *P* - value* |
| --- | --- | --- |
| Gpg1A vs. Gpg2A | 445,282 vs. 513,094 | 0.596 |
| Gpg1C vs. Gpg2C | 178,020 vs. 235,183 | 0.371 |
| Gpg1H vs. Gpg2H | 171,843 vs. 167,841 | 0.884 |
| Gpg1A vs. Gpg1C | 445,282 vs. 178,020 | $4.34 \times 10^{-3}$ |
| Gpg1A vs. Gpg1H | 445,282 vs. 171,843 | $1.86 \times 10^{-3}$ |
| Gpg1C vs. Gpg1H | 178,020 vs. 171,843 | 0.862 |
| Gpg2A vs. Gpg2C | 513,094 vs. 235,183 | $1.38 \times 10^{-2}$ |
| Gpg2A vs. Gpg2H | 513,094 vs. 167,841 | $6.92 \times 10^{-4}$ |
| Gpg2C vs. Gpg2H | 235,183 vs. 167,841 | 0.295 |

\**p* values were calculated using the Negative Binomial Regression model after adjusting for covariance.

A total of 11,398 microbial species was identified in the 161 dental plaque samples using sequence or phylogenetic similarity to microNR. An average of more than 3,000 species were predicted in each dental plaque sample from AAs, which was significantly higher than those found in the samples from CAs and HAs (Table 3). In addition, distinctive microbial species were identified in AAs, CAs, and HAs, which is consistent with findings of unique genes among the groups. There were 501 microbial species found only in the AA group compared to 153 and 310 detected in its counterparts of CAs and HAs, respectively (Fig. 2). Table 4 presents the race/ethnic specific-species that were detected in 5 or more samples of each particular racial/ethnic group. Interestingly, one bacterial species, *Pedobacter petrophilus*, was only detected in the samples from AAs with higher level of *P. gingivalis* (Gpg2A). These results imply that specific microbial profiles contribute to periodontal health disparities.

**Fig. 2.**
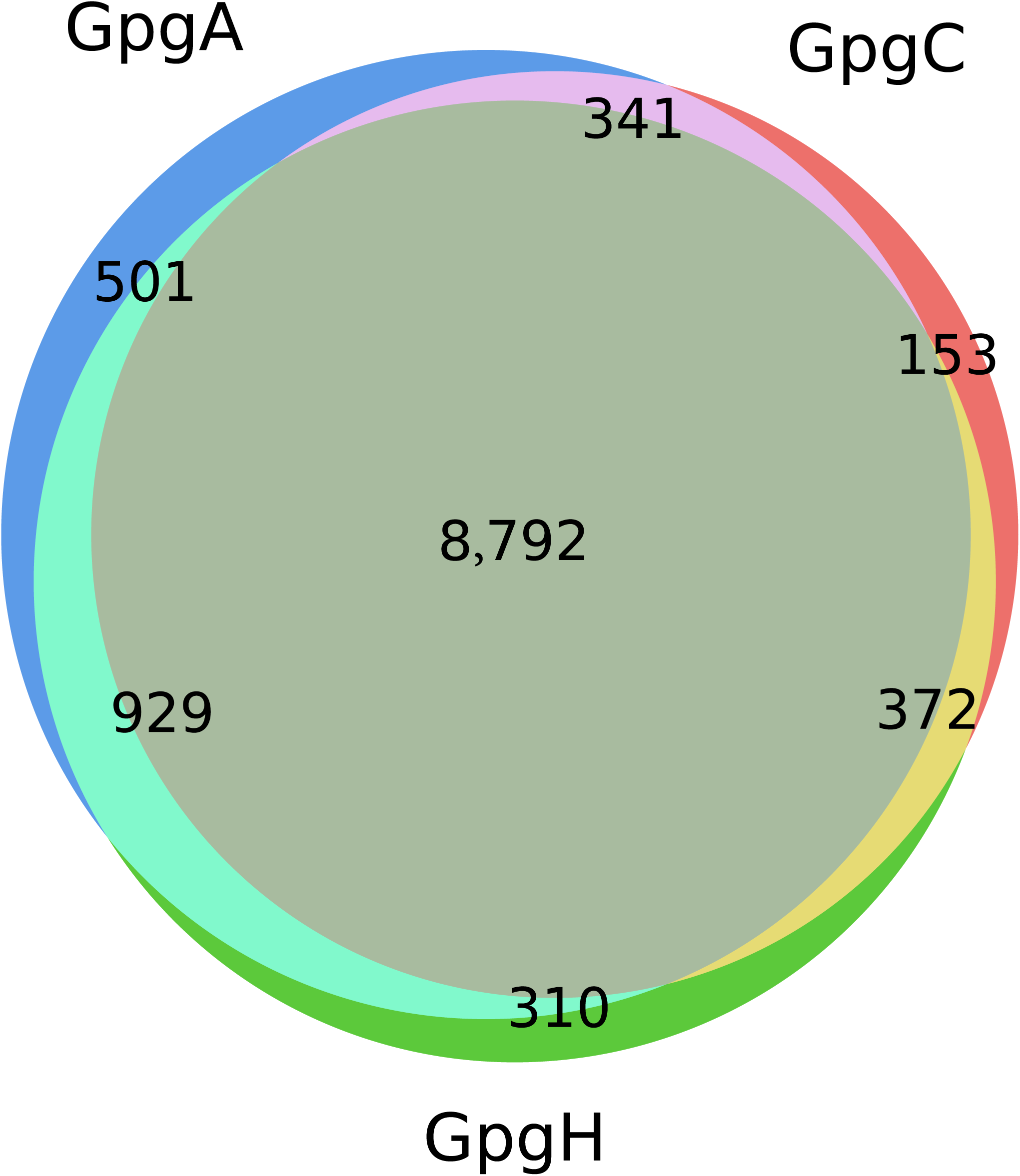
Venn diagram of the microbial taxa at the species level identified in the racial/ethnic groups AAs (GpgA), CAs (GpgC), HAs (GpgH).

**Table 3.** The mean number of taxonomies identified per samples in different groups.

| Comparison of<br>sample groups | Mean number of<br>species | *P* - value* |
| --- | --- | --- |
| Gpg1A vs. Gpg2A | 3,184 vs. 3,893 | 0.299 |
| Gpg1C vs. Gpg2C | 2,028 vs. 2,268 | 0.165 |
| Gpg1H vs. Gpg2H | 1,836 vs. 2,030 | 0.349 |
| Gpg1A vs. Gpg1C | 3,184 vs. 2,028 | $2.00 \times 10^{-3}$ |
| Gpg1A vs. Gpg1H | 3,184 vs. 1,836 | $4.19 \times 10^{-4}$ |
| Gpg1C vs. Gpg1H | 2,028 vs. 1,836 | 0.860 |
| Gpg2A vs. Gpg2C | 3,893 vs. 2,268 | $7.17 \times 10^{-3}$ |
| Gpg2A vs. Gpg2H | 3,893 vs. 2,030 | $8.26 \times 10^{-5}$ |
| Gpg2C vs. Gpg2H | 2,268 vs. 2,030 | 0.172 |

\**P* values were calculated using the Negative Binomial Regression model after adjusting for covariance.

**Table 4.** Unique bacterial species found in AAs and HAs.

| Numbers of samples with the unique species |  |  |  |  |  |
| --- | --- | --- | --- | --- | --- |
| AAs |  |  | HAs |  |  |
| Species | Gpg1A | Gpg2A* | Species | Gpg1H | Gpg2H |
| *Streptomyces sp. WAC04114* | 2 | 3 | *Micrococcales bacterium* | 3 | 5 |
| *Vagococcus hydrophili* | 5 | 0 | *Megasphaera sp. DISK 18* | 2 | 4 |
| *Veillonella sp. OK1* | 4 | 2 | *Ruminococcus sp. NSJ-71* | 3 | 3 |
| *Pedobacter petrophilus* | 0 | 7 | *Ideonella azotifigens* | 1 | 4 |
| *Hyphomicrobium sp. CS1GBMeth3* | 1 | 5 | *Pseudomonas caricapapayae* | 4 | 2 |
| *Halomonas zhangzhouensis* | 3 | 2 | *Siphovirus Jomon_CT89* | 2 | 4 |

\*Gpg1 includes samples with the median abundance (FPKM) of *P.*
*gingivalis* (<5,741 FPKM), and Gpg2 had samples with the median abundance (FPKM) of *P. gingivalis* (>59,862 FPKM).

Notably, *P. gingivalis* were identified in all 161 samples with whole metagenome shotgun sequencing even though *P. gingivalis* was detected in only 50% samples in this cohort using qPCR. However, samples (Gpg 2) in which *P. gingivalis* can be detected using PCR had ten-fold higher abundance of *P. gingivalis* than those (Gpg 1) with *P. gingivalis* not detected using PCR (Table 5). Therefore, we re-designated Gpg 1 as the group with low level of *P. gingivalis* and Gpg 2 as the high-level *P. gingivalis* group. In addition, *Tannerella forsythia*, *Fusobacterium nucleatum*, *Streptococcus cristatus*, and *Streptococcus gordonii* were also detected in all 161 samples. *Treponema denticola* and *Filifactor alocis* were identified in 160 and 158 samples out of 161 samples, respectively. We further determined the abundance of several well-studied bacterial species. We observed significantly more abundance of *P. gingivalis* and *T. denticola* in the dental samples retrieved from AA and HA groups compared to CA group (Table 6). Whereas, *F. alocis* and *T. forsythia* were detected significantly more in HAs than CAs. We did not observe any significant difference in abundance of *F. nucleatum* and *S. cristatus* among three groups. we also found a significantly higher number of S. gordonii in AAs than that in HAs, but not between AAs and CAs.

**Table 5.** Detection rates of well-known oral bacteria in different groups.

| Sample group | Gpg1A | Gpg1C | Gpg1H | Gpg2A | Gpg2C | Gpg2H | Gpg1 | Gpg2 |
| --- | --- | --- | --- | --- | --- | --- | --- | --- |
| Samples count | 22 | 21 | 38 | 22 | 21 | 37 | 81 | 80 |
| *P. gingivalis* | 22 | 21 | 38 | 22 | 21 | 37 | 81 | 80 |
| *F. alocis* | 22 | 20 | 37 | 22 | 21 | 36 | 79 | 79 |
| *T. forsythia* | 22 | 21 | 38 | 22 | 21 | 37 | 81 | 80 |
| *T. denticola* | 22 | 21 | 37 | 22 | 21 | 37 | 80 | 80 |
| *F. nucleatum* | 22 | 21 | 38 | 22 | 21 | 37 | 81 | 80 |
| *S. cristatus* | 22 | 21 | 38 | 22 | 21 | 37 | 81 | 80 |
| *S. gordonii* | 22 | 21 | 38 | 22 | 21 | 37 | 81 | 80 |

**Table 6.** The average abundance (FPKM) of species per sample in different groups.

| Sample group | GpgA | GpgC | *p*-value | GpgA | GpgH | *p*-value | GpgC | GpgH | *p*-value* |
| --- | --- | --- | --- | --- | --- | --- | --- | --- | --- |
| *Porphyromonas gingivalis* | 159,821 | 82,068 | 0.002 | 159,821 | 112,880 | 0.015 | 82,068 | 112,880 | 0.021 |
| *Filifactor alocis* | 48,664 | 28,642 | 0.086 | 48,664 | 22,435 | 0.102 | 28,642 | 22,435 | 0.007 |
| *Tannerella forsythia* | 102,822 | 73,717 | 0.133 | 102,822 | 104,979 | 0.441 | 73,717 | 104,979 | 0.039 |
| *Treponema denticola* | 64,395 | 34,671 | 0.001 | 64,395 | 83,108 | 0.115 | 34,671 | 83,108 | 0.0003 |
| *Fusobacterium nucleatum* | 323,709 | 233,939 | 0.392 | 323,709 | 189,829 | 0.254 | 233,939 | 189,829 | 0.860 |
| *Streptococcus cristatus* | 26,989 | 28,667 | 0.749 | 26,989 | 29,825 | 0.908 | 28,667 | 29,825 | 0.712 |
| *Streptococcus gordonii* | 38,290 | 29,472 | 0.110 | 38,290 | 23,824 | 0.005 | 29,472 | 23,824 | 0.199 |

\**P* values were calculated by fitting a linear model (adjusting for covariance) and then applying Kruskal-Wallis test to the residuals extracted from the model for the racial groups under comparison.

To identify potential patterns of oral microbial profiles in different racial/ethnic groups, we projected the dental plaque samples from high-dimensional space into a two-dimensional space using NMDS, a non-supervised machine learning technique. NMDS retains the original sample rank-order similarity by measuring it using Euclidean distance between the points that represent the samples. The visualization of the NMDS result is provided in Fig. 3, which shows that the AA groups, Gpg1A and Gpg2A, are both located in the leftmost section of the figure and are much closer to one another than CAs and HAs (Fig. 3), indicating that the microbial profiles are more similar among the samples in the AA groups than those of the other two groups. Additionally, both Gpg1H and Gpg2H, i.e., the HA groups, were projected toward the bottom right corner of the figure, further away from the AA groups than the EA groups. More interestingly, the Gpg1 groups (including Gpg1A, Gpg1C, and Gpg1H) occupy the upper section of the figure more than the Gpg2 groups (i.e., Gpg2A, Gpg2C, and Gpg2H), showing distinction between the two groups which were determined based on the abundance of *P. gingivalis*.

**Fig. 3.**
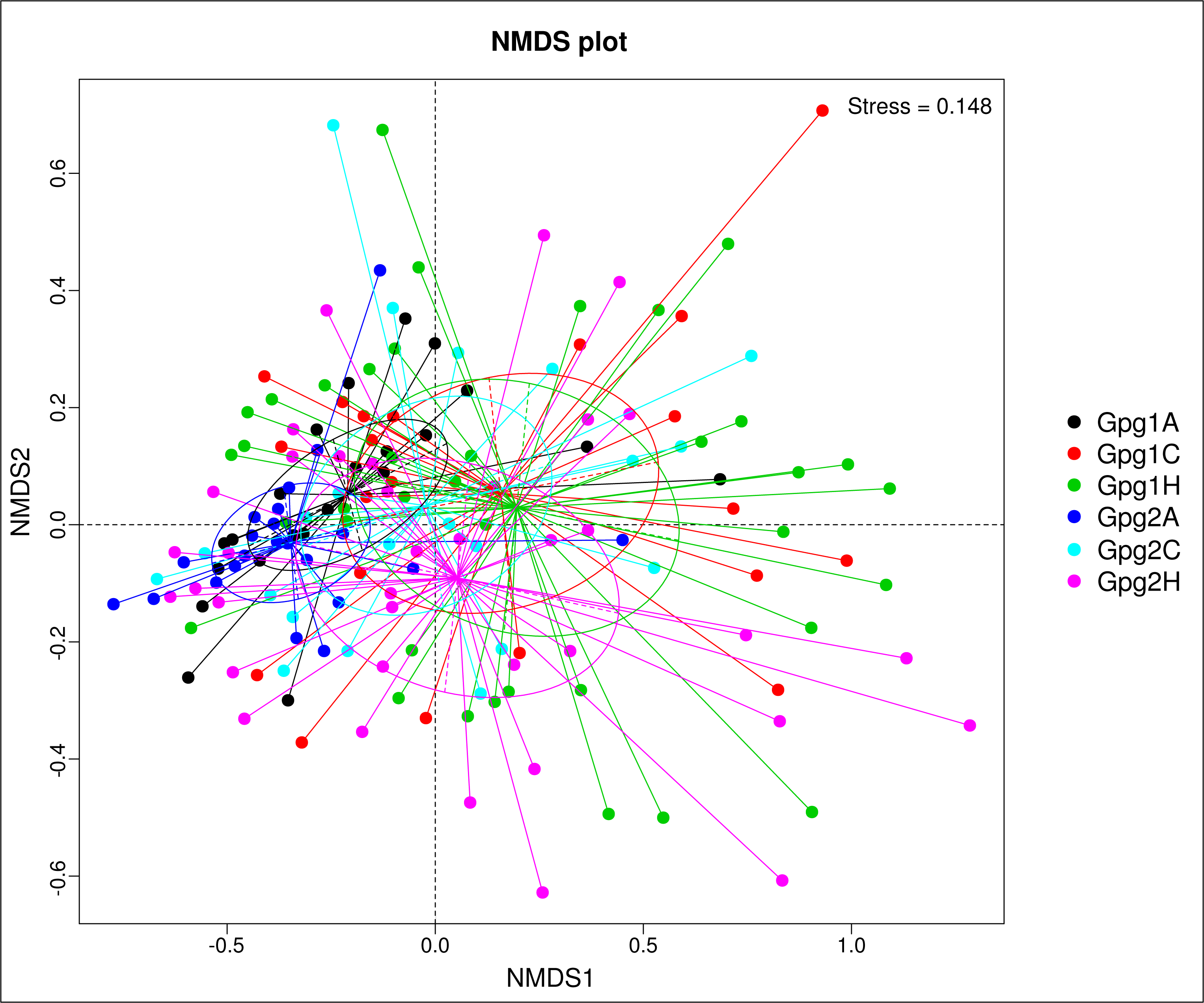
Visualization of NMDS analysis. Dots in a two-dimensional space represent dental plaque samples and the distance between each pair of dots represents the dissimilarity between the corresponding two samples. The samples in the same groups were assigned the same colors.

### Functional diversities of the oral microbiome associated with levels of *P. gingivalis* and racial/ethnic backgrounds

Functional annotation was conducted by assembling metagenomic sequencing and mapping against functional protein databases. Mapping against the Comprehensive Antibiotic Resistance Database (CARD) allowed us to identified a total of 68 antibiotic resistance genes (ARGs) in the 161 dental plaque samples (Fig. 4A). The median numbers of ARGs per sample were 40 in the plaque samples from the AA group, 34 ARGs from the CAs, and 28 ARGs from the HAs. Additionally, more abundant ARGs (approximate medium copies of 12,600) were also detected in the samples from the AAs, compared to 10,600 and 9,500 in the samples from the CAs and the HAs, respectively (Fig. 4B). These data demonstrate more complex and abundant ARGs were present in the oral microbiome of AAs compared to those in CAs and HAs, suggesting that the oral microbiomes of AAs may have a higher ability to resist antibiotics. Moreover, Fig. 5 presents the functions of the ARGs. The majority of ARGs act on antibiotic efflux and inactivation, and others are involved in antibiotic target alteration, protection, and replacement. Most of the ARGs were found in *Bacillota*, a phylum of mostly Gram-positive bacteria, *Pseudomonadota* including Gram-positive, negative, and variable bacterial species, and *Bacteroidota*, a phylum of Gram-negative bacteria (Fig. 5), which presumably serve as reservoirs for ARGs in the oral microbiome.

**Fig. 4.**
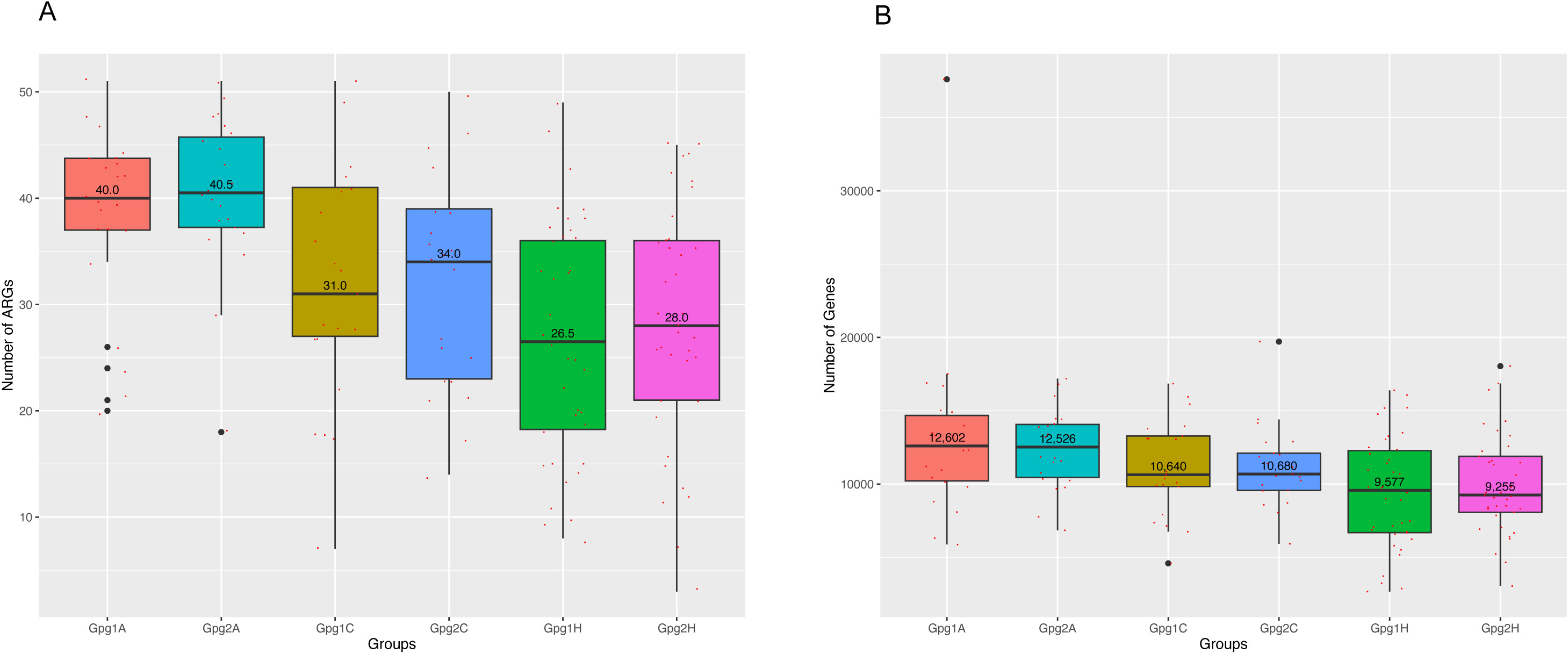
Diversities and abundances of antibiotic-resistant genes (ARG). (A) Total number of ARG classes, and (B) The abundances of ARGs in each sample. Each sample is presented as a red point and the number in each boxplot represents median number of ARG classes and abundances in a group.

**Fig. 5.**
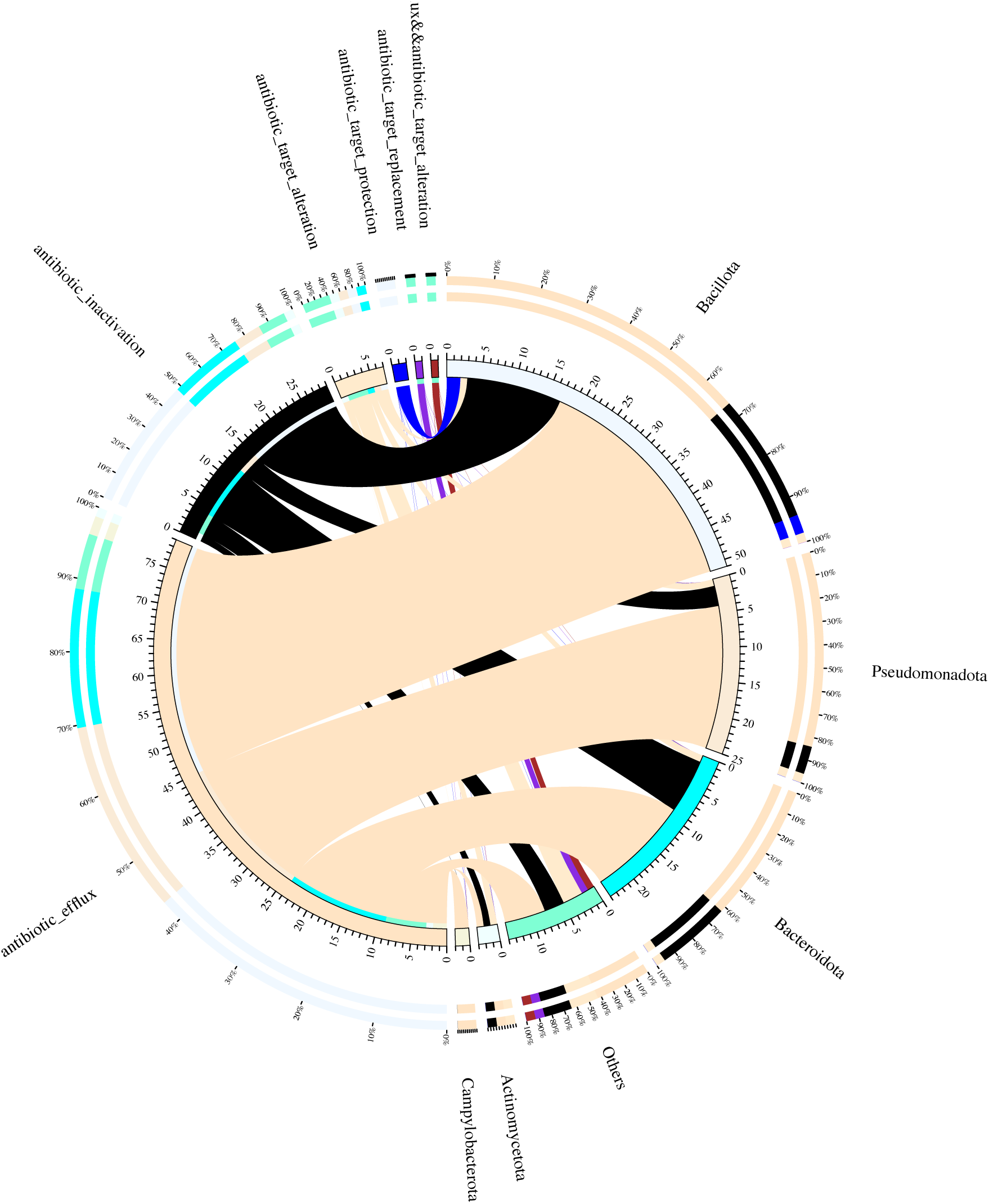
Antibiotic resistance genes mechanism and the loop graph of species distribution. Circle chart is divided into two parts, the right side shows the sample information, the left side shows the ARG tolerance of antibiotic information. Inner circle different colors represent different samples and ARGs, scale for the relative abundance (unit ppm). The left side is the sum of the relative abundance of the resistance genes in the sample, the right side is the sum of the relative abundance of the resistance genes in each ARG. The left side of the outer circle shows the relative percentage of the antibiotic to which the resistance gene belongs to, and the right side of the outer ring shows the relative percentage of the sample in which the antibiotic resistance gene is located.

To determine the physiology of racial/ethnic-associated oral microbiome, we also analyzed the assembled metagenomic protein sequences against the Carbohydrate-Active enZYmes Database (CAZy) (18). A cluster heatmap shows the quantitation of the top 35 carbohydrate-active enzymes involving in the breakdown, biosynthesis, and modification of glycoconjugates, oligo- and polysaccharides (LPS). The genes encoding these enzymes were much more abundant in the oral microbiome of the AAs, compared to those in the CAs and HAs (Fig. 6). This implies that Gram-negative bacteria in the oral microbial communities may be relatively higher in AA population than in CAs and HAs, which can induce immune-inflammatory responses and initiates the onset of periodontitis.

**Fig. 6.**
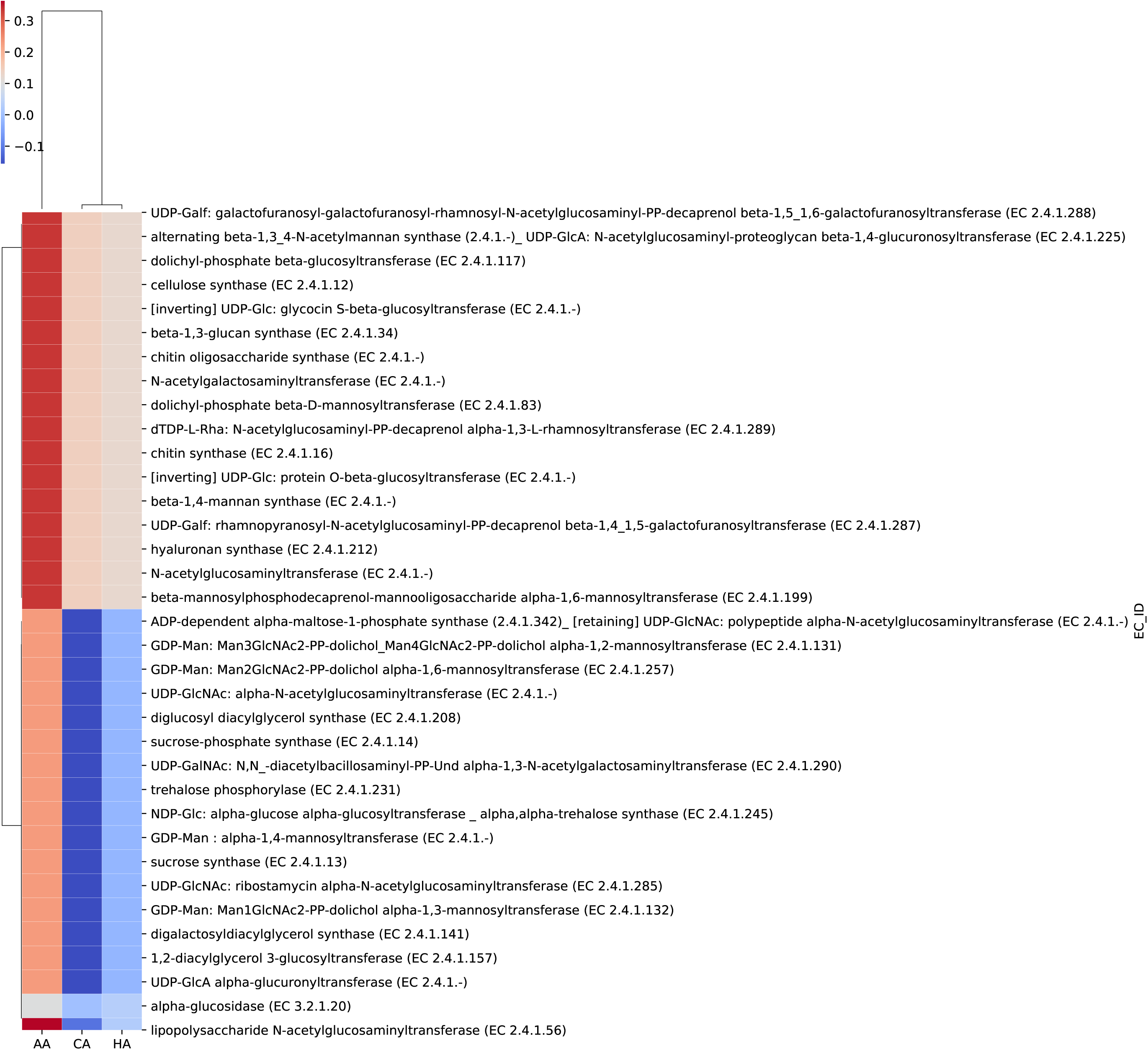
Heatmap of carbohydrate-active enzyme genes identified in AA, CA, and HA groups. Columns represent sample groups AA, CA, and HA and rows represent genes. The red color represents high gene abundance to contrast with low abundance in blue color. The rows and columns are ordered based on the correlations of z scores which were calculated based on gene abundance.

## DISCUSSION

In this paper, we investigated variations of the oral microbial communities across races and ethnicities using whole metagenomic sequencing. Recent studies of the oral microbiome, which mainly used less powerful qPCR and 16S rRNA gene sequencing, found significant difference in microbial compositions between/among racial/ethnic populations, including higher abundance of *P. gingivalis*, *T. forsythia*, *T. denticola*, and *F. alocis* in AAs (15, 19–21). Our results on the cohort with intact periodontium are consistent with the previous reports that some well-known periodontal pathogens including *P. gingivalis* and *T. forsythia* were more abundant in AAs and HAs than those in CAs. However, comparing to previous qPCR-based studies that only detected *P. gingivalis* at approximately 25% of samples from individuals without periodontitis (17, 22), we found *P. gingivalis* in all tested dental plaques in this study with whole metagenomic sequencing, even though we previously did not identified the *P. gingivalis* in half of the samples using qPCR. Moreover, *T. forsythia* was also found in all samples, while detection rates of *T. denticola* and *F. alocis* were 99% and 98%, respectively. These results are not in agreement with some other studies that reported much lower detection rates of these bacteria using qPCR. For example, *T. forsythia* was detected in 10.5% of periodontally healthy participants (32) and 68.0% of periodontitis patients (33). Moreover, our findings indicate that there is no difference in the detection rates of most well-known periodontal pathogens among racial/ethnic groups. These results indicate whole metagenomic sequencing is more sensitive than qPCR and 16S rRNA Sequencing for microbial detection at the species level of all taxa including viruses. Therefore, detection rates of periodontal pathogens may not be closely associated with periodontal health disparities, rather it is the levels of these pathogens in the oral microbiome that contribute to the initiation and development of periodontitis. In addition, AAs, CAs, and HAs differ in microbial diversities. We identified higher numbers of microbial taxa in the oral microbiome of AAs and HAs than CAs and greater diversities of microbial compositions in AAs and HAs than CAs. Racial/ethnic associated microbial species has not been reported to be associated with any human oral disease so far. Interestingly, we identified that some bacterial species were only found in the oral microbiota of AAs, such as *P. petrophilus,* which was clearly associated with higher levels of *P. gingivalis*. This bacterial species is currently not easily cultured in laboratory, and hence difficult to analyze its relationship with other well-known periodontitis pathogens and its role in the initiating periodontitis. Nevertheless, the unique presentence of this bacterial species in the oral cavities of AA population may lead to a new research pathway linking to periodontal health disparities.

One advantage of whole metagenomic sequencing is its ability to reveal information on functional potentials of the oral microbiome, enabling us to compare antibiotic resistance and carbohydrate enzyme activities of the oral microbiome among AAs, CAs, and CAs. Using metagenomic sequencing, we demonstrated more diverse and abundant ARGs in the oral microbiome of the AA group, which may be an important determinant of virulence that influence the ability of microorganisms to adapt and survive in the oral cavities. It remains to be determined how complex and abundant ARGs evolve in the oral microbiome of the AA population. One possible mechanism is that inappropriate antibiotic use was more common among AA patients compared to CA patients (23). Another unique characteristic of the oral microbiome of AAs is its higher level of enzymes involving in modification of glycoconjugates, oligo- and polysaccharides, which may be linked to higher level of LPS. This observation is consistent with the levels of several Gram-negative periodontal pathogens including *P. gingivalis*, *T. forsythia*, *T. denticola*, and *F. alocis*.

Besides identification of functional potentials, other strengths of our study include a comprehensive periodontal examination for each participant and taxonomic annotation coverage at every taxonomic level including the species level. A previous study, using 16s rRNA sequencing, reported that African American had higher microbial diversity in oral washes than Caucasians (15). By contrast, another study indicated that African American adults had the lowest bacterial diversity in subgingival plaques while Chinese and Caucasian adults had the highest diversity (24). Both of these studies lack detailed oral health parameters and used 16s rRNA sequencing. It should be pointed out that one limitation of this study is based on self-reported race and ethnicity data. However, previous studies suggested self-identified race and ethnicity of AAs and CAs are generally reliable (25). In the present study, we observed significant difference in the oral microbiome among AA and CA populations, and relatively less difference found between CAs and HAs, which may be due to HA population with more complex genetic makeup. More accurate information on ancestry proportions of the participants will be included in HAs for future health disparity studies.

In summary, this study shows significant differences in oral microbial diversity and abundance among AA, CA, and HA groups. We also identified racial/ethnic specific bacterial species, such as *P. petrophilus* that was only found in AAs with relatively higher *P. gingivalis* levels. Moreover, higher functional potential including antibiotic resistance and LPS production were observed in the oral microbiome of AAs. These differences may shape virulence potential of the oral microbiome and prepare microbiome as a whole to adapt their environmental niches. Understanding and comparing racial/ethnic associated oral microbiome provide a steppingstone for improving oral health disparities.

## MATERIALS AND METHODS

### Study cohorts

The research protocol was approved by the Committee for the Protection of Human Subjects of the University of Texas Health Science Center at Houston (IRB number: HSC-DB-17-0636). Candidates were screened during routine dental visits at the clinic of the School of Dentistry, University of Texas Health Science Center at Houston between 2017 and 2022. Individuals aged 21-75 years with self-reported ethnicity/race of AAs, CAs, or HAs were enrolled after the initial periodontal examination that included determination of plaque index (PI), bleeding on probing (BOP) level, probing depth, and clinical attachment level on all teeth (**26**). Radiographs were taken during this screening phase to assess bone loss. The clinical oral examinations were performed by faculty members of the School of Dentistry, University of Texas Health Science Center at Houston. The examiners are calibrated annually in the diagnosis of periodontitis. Based on the 2017 World Workshop classification (27, 28), all study participants diagnosed as clinical gingival health or biofilm-induced gingivitis on an intact periodontium met the following criteria: >24 teeth; no alveolar bone loss or clinical attachment loss; pocket depth ≤3mm (excluding pseudo pocket); no antibiotic therapy in the previous six months; and not pregnant.

### Dental plaque sample collection

Dental plaque samples, including supra and subgingival dental plaques, were collected by board-certified periodontists using sterile paper points prior to any dental treatment and labelled numerically according to the sampling sequences. The paper points were placed in the sulci of all the first molar in different quadrants for 1 min and then immersed immediately in an Eppendorf tube with 0.5 ml of Tris-EDTA (TE) buffer (pH 7.5) (29). Bacterial pellets were harvested by centrifugation and then resuspended in 100 µl TE buffer. Samples were stored at −80°C until use.

### Sequencing and quality control

Samples were sent to Novogene Co. (Sacramento, CA, USA) with dry ice for metagenomic sequencing. Briefly, DNA extracted from human dental plaques was randomly sheared into short fragments, and the resulting fragments were end-repaired, A-tailed, and ligated using Illumina adapters. The fragments with adapters were subsequently amplified using PCR, followed by size selected and purified. Quality control of the library was conducted using Qubit (≥20 ng, ≥10ng/μl), and quantification and size distribution detection were performed using real-time and a bioanalyzer, respectively. The quantified libraries were pooled and sequenced using an Illumina NovaSeq high throughput sequencer by Novogene Corporation, Inc., with a paired-end sequencing length of 150 bp and an output of ∼6 GB of raw data per sample.

### Data preprocessing

The average size of raw data generated per sample was 6.394 GB. To ensure accuracy and reliability of the subsequent data analysis, all low-quality bases (Q-value ≤ 38) that exceed certain threshold (40 bp), as well as reads containing N nucleotides over 10 bp and those overlapping with adapters > 15 bp were trimmed. After quality control, the size of clean data was 6.393 GB per sample, with 96.36% and 91.26% of bases having quality scores greater than 20 and 30, respectively. To minimize host DNA contamination, raw reads that were mapped to the human reference genome were discarded using the Bowtie2 software (30).

### Gene prediction and abundance analysis

The MetaGeneMark software was used to predict open reading frames (ORFs) from scaftigs (> = 500bp) (31). ORFs less than 100 nt were discarded. To generate gene catalogues, the remaining ORFs were dereplicated using CD-HIT (32, 33) in default settings (i.e., identity = 95 %, coverage = 90 %). To calculate the gene quantity, the clean data were mapped to the gene catalogue using Bowtie2 (parameters: –end-to-end, –sensitive, –I 200, –X 400). Gene abundance (*G_k_*) was calculated using the following formula:

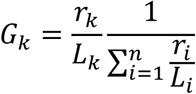

where *r* represents the number of mapped reads and *L* represents gene length. Downstream analyses were performed based on the abundance of the gene catalogues.

### Taxonomy annotation

The software tool DIAMOND (version 0.9.9.110) (34) was used to align the sequences of the identified genes to those of bacteria, fungi, archaea, and viruses extracted from NCBI’s NR database (version 2018-01-02). MEGAN software was used to taxonomically annotate each metagenomic homolog (35). The sum of abundance of genes annotated as a species in a sample was used as the abundance estimate of that species in that sample. Based on the abundance of each taxonomic level, various analyses were performed, including heatmap of abundance, principal coordinate analysis (PCoA), principal component analysis (PCA), and non-metric multi-dimensional scaling (NMDS) analysis, which is an indirect gradient analysis approach that produces ordination based on a distance matrix. R package ade4 (version 3.2.1) was used to perform PCA analysis and R package vegan (version 2.15.3) was used for PCoA and NMDS analyses.

### Functional analysis

The identified gene sequences were aligned to those in functional databases utilizing the DIAMOND software (version 0.9.9.110) (34), with parameter settings: blastp, -e 1e-5. The functional databases used in this study include Comprehensive Antibiotic Resistance Database (CARD) (36), Carbohydrate-Active Enzymes Database (CAZy) (18), Kyoto Encyclopedia of Genes and Genomes (KEGG) (37), and Genealogy of Genes: Non-supervised Orthologous Groups (eggNOG) (38). Based on the sequence alignment results, the best Blast hits were selected for subsequent analysis and the relative abundance were calculated at different functional levels.

### Statistical analysis

Statistical analyses were conducted using statsmodels (version 0.10.1) and SciPy (version 1.4.1), open-source Python libraries for statistical modelling and scientific computing. Chi-square tests were performed to compare gender of different racial/ethnic groups. Kruskal-Wallis tests were applied to compare ages of the sample groups. For the total number of unique genes and taxa characterized in each sample, Negative Binomial Regression was performed to adjust for age and gender while comparing the three racial groups. In addition, to compare the abundance of a particular species in the three racial groups, linear regression was firstly performed to fit the data, adjusting for covariates (age and gender), Kruskal-Wallis test was then applied to the residuals extracted from the regression model to calculate p values. A *p*-value of < 0.05 was considered statistically significant.

## ACKNOWLEDGMENTS

The authors are grateful to all study participants for their contributions to this research. The authors thank Krishna Kookal for abstracting clinical parameters from the Electronic Health Record at the School of Dentistry, University of Texas Health Science Center at Houston.

The study was supported in part by grant U54MD007586 from the National Institute on Minority Health and Health Disparities (NIMHD), USA; grant R16GM149359 from the National Institute of General Medical Sciences (NIGMS), USA; grant UG3HG013248 from the National Human Genome Research Institute (NHGRI), USA; and grant 1OT2OD032581 from the National Institutes of Health (NIH), USA. The work is solely the responsibility of the authors and should not be interpreted as representing the official policies, either expressed or implied, of the NIH.

H.X. contributed conception and design of the study. Q.W. and H.X. performed data analysis and prepared Tables and Figures. B.W. prepared Table 1 and collected dental plaque samples. All authors were involved in manuscript writing and revision.

## ETHICS APPROVAL

The research protocol was approved by the Committee for the Protection of Human Subjects of the University of Texas Health Science Center at Houston (IRB number: HSC-DB-17-0636). Consent forms were signed by all participants, before sample collection.

## DATA AVAILABILITY

Our metagenomic sequencing data are deposited at Sequence Read Archive (SRA) with accession number PRJNA1160290.

## PATIENT CONSENT FOR PUBLICATION

We give our consent for the publication of the manuscript including all tables and figures.

## CONFLICT OF INTEREST

All authors declare no potential conflicts of interest with respect to the research, authorship, and/or publication of this article.

